## Supplementary Figures and Table 1,2,5,7-11,19 for "Biobank-wide association scan identifies risk factors for late-onset Alzheimer’s disease and endophenotypes"

**Supplementary Figure 1.** **A** **flowchart for analyses of Alzheimer’s genetic data.**

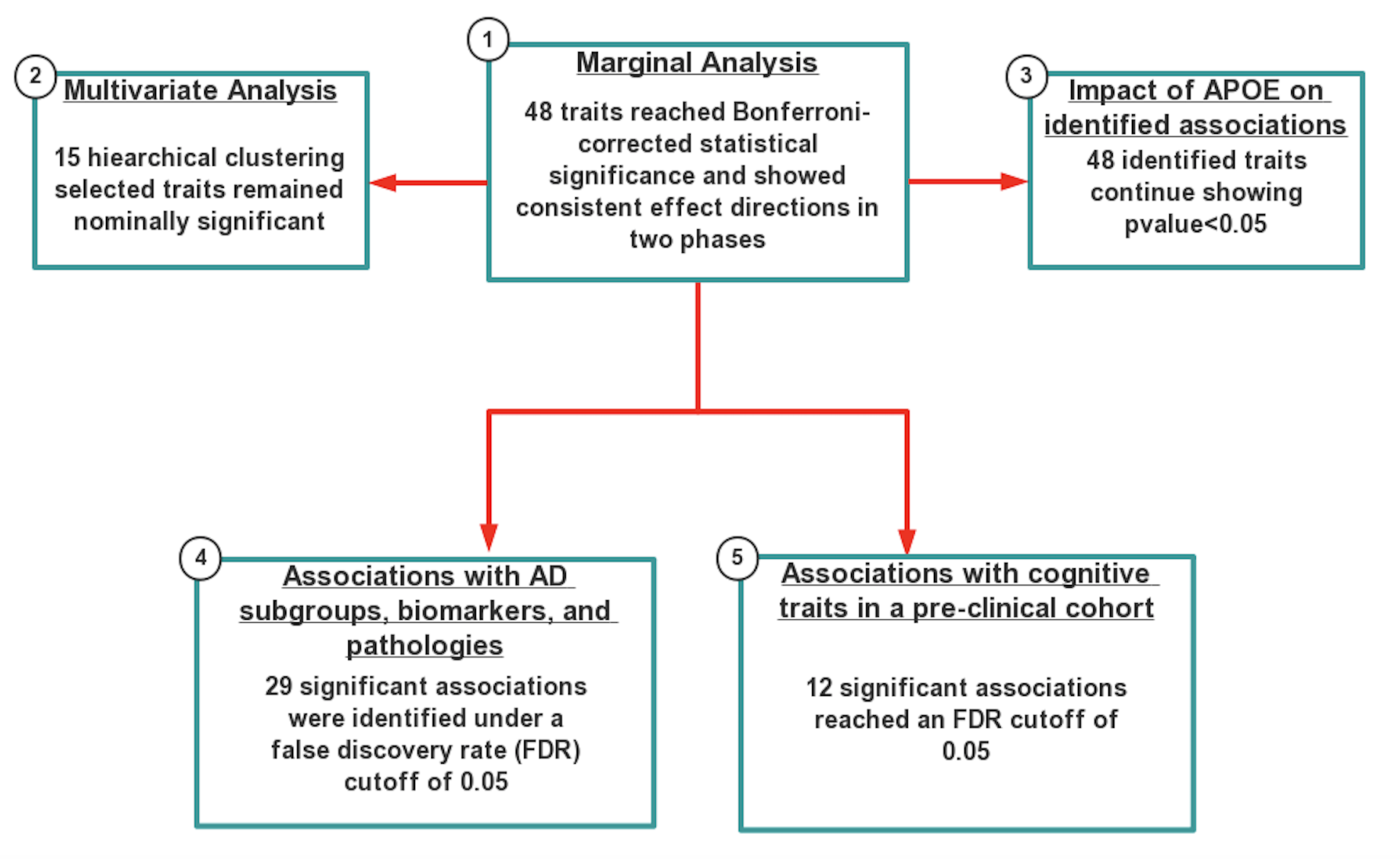

**Supplementary Figure 2. Comparison of effect size estimates from BADGERS and regression analysis based on individual-level data.** BADGERS and regression analysis showed highly consistent effect size estimates for 1,738 PRS in simulation setting 2.

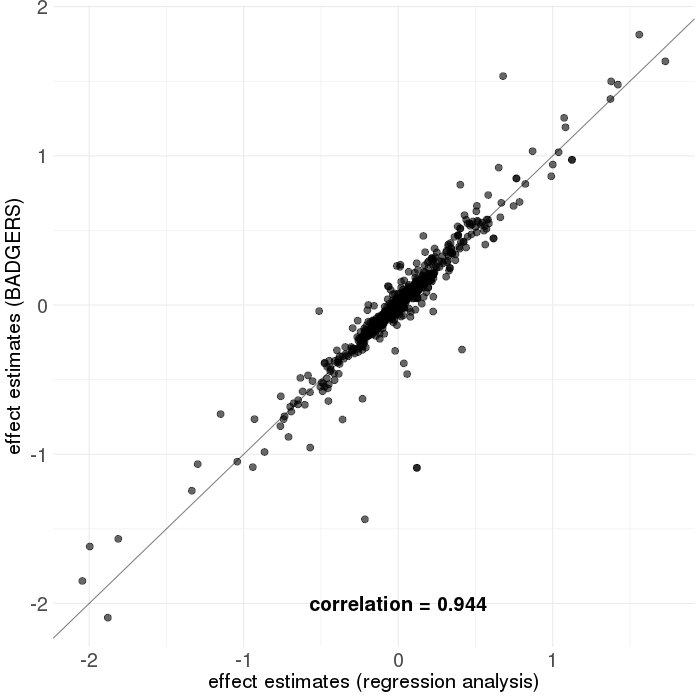

**Supplementary Figure 3. Comparison of p-values from BADGERS and regression analysis based on individual-level data.** BADGERS and regression analysis provided highly consistent p-value results for 1,738 PRS in simulation **(A)** setting 1 and **(B)** setting 2.

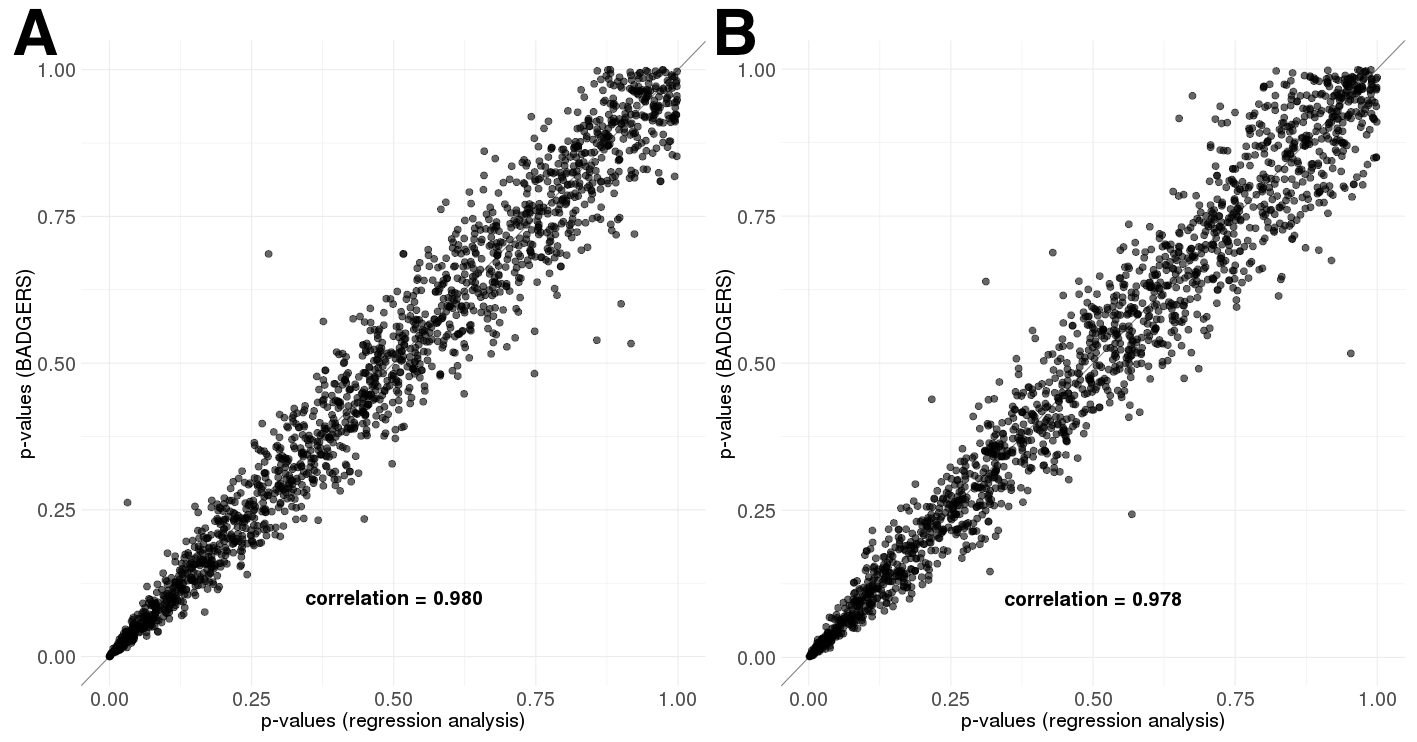

**Supplementary Figure 4.** **Comparison of effect size estimates from BADGERS and regression analysis based on individual-level data when p-values are smaller than 0.05.** The effect size between two algorithms is highly consistent.

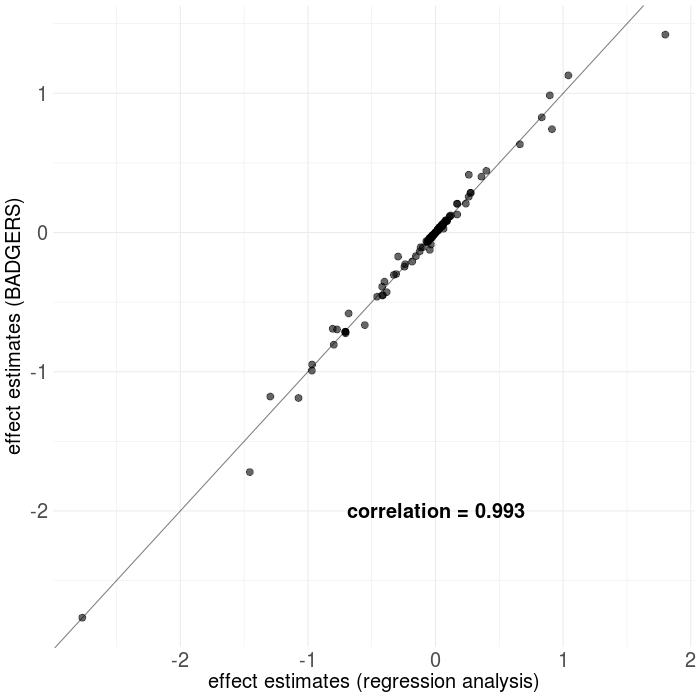

**Supplementary Figure 5. BADGERS estimates using marginal PRS and joint PRS.** Both methods showed consistent effect size estimates with true effect size in simulation.

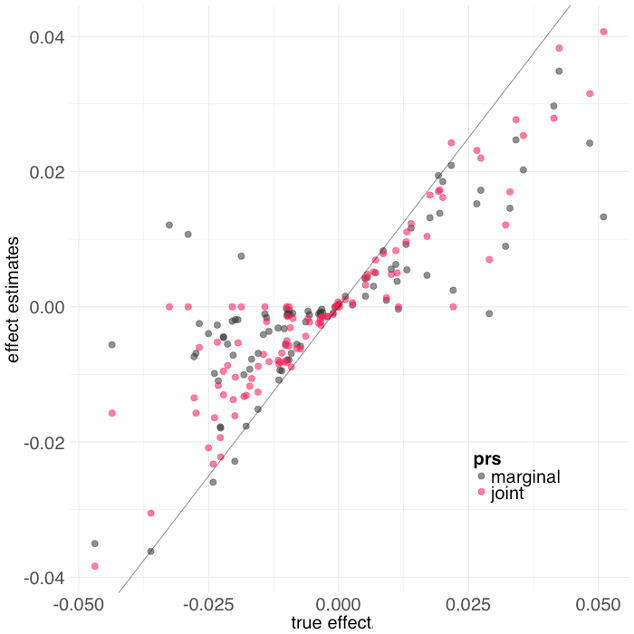

**Supplementary Figure 6. Workflow of the two-stage BWAS for late-onset AD.**

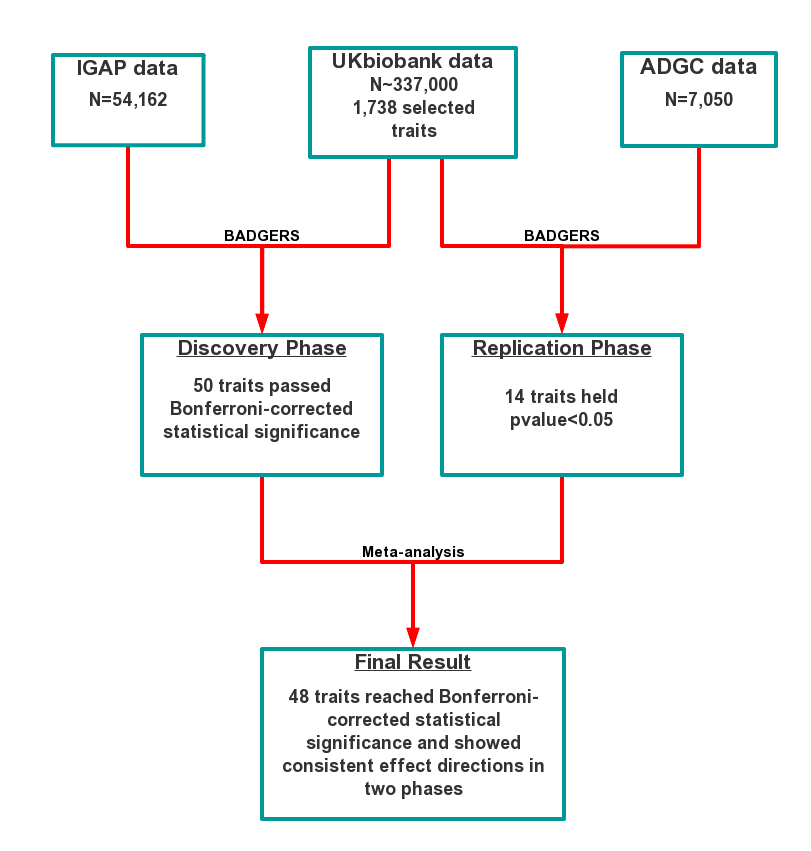

**Supplementary Figure 7. Associations between AD and education attainment in two independent analyses.** Error bars denote the standard error of effect estimates.

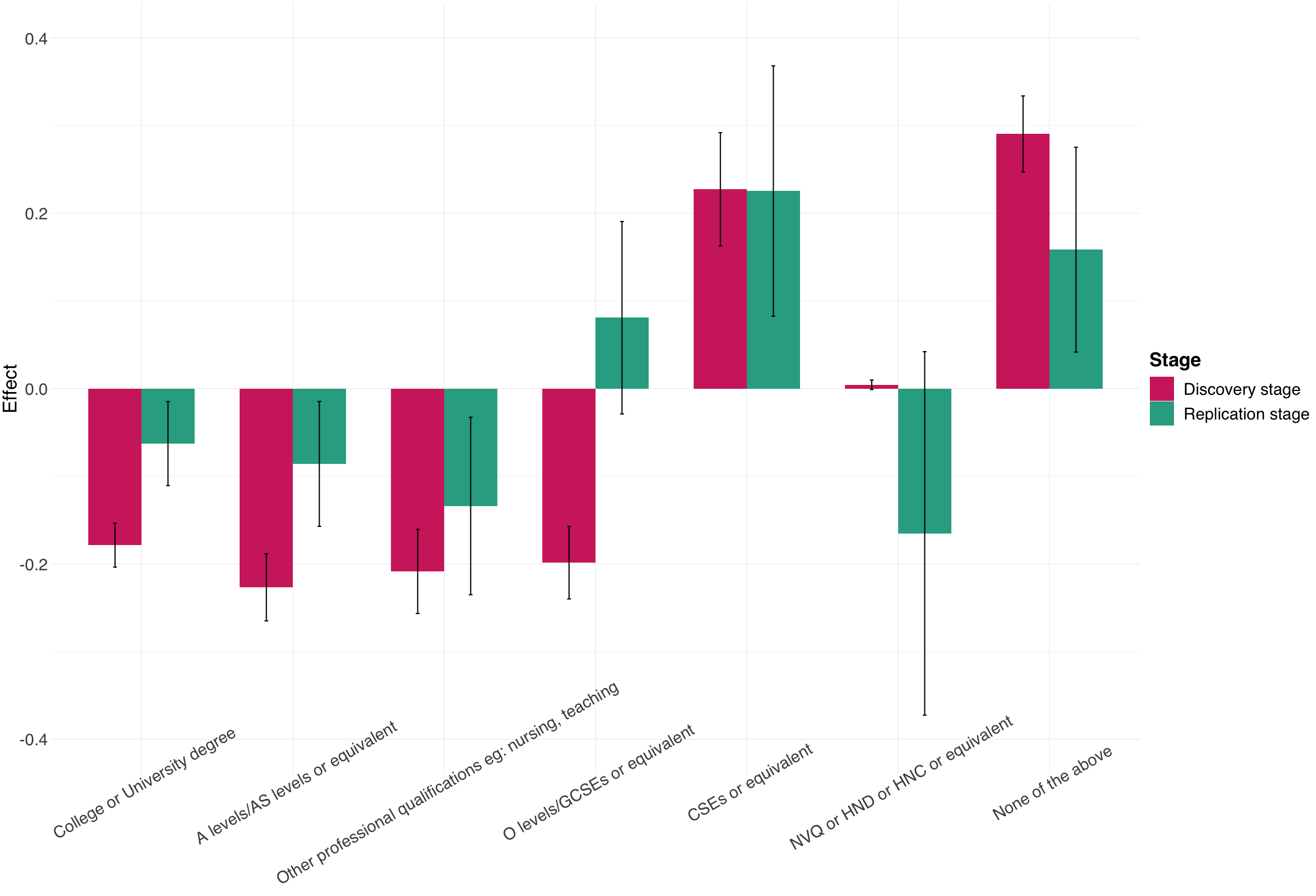

**Supplementary Figure 8. Correlation heatmap for the 15 representative traits selected based on hierarchical clustering.**

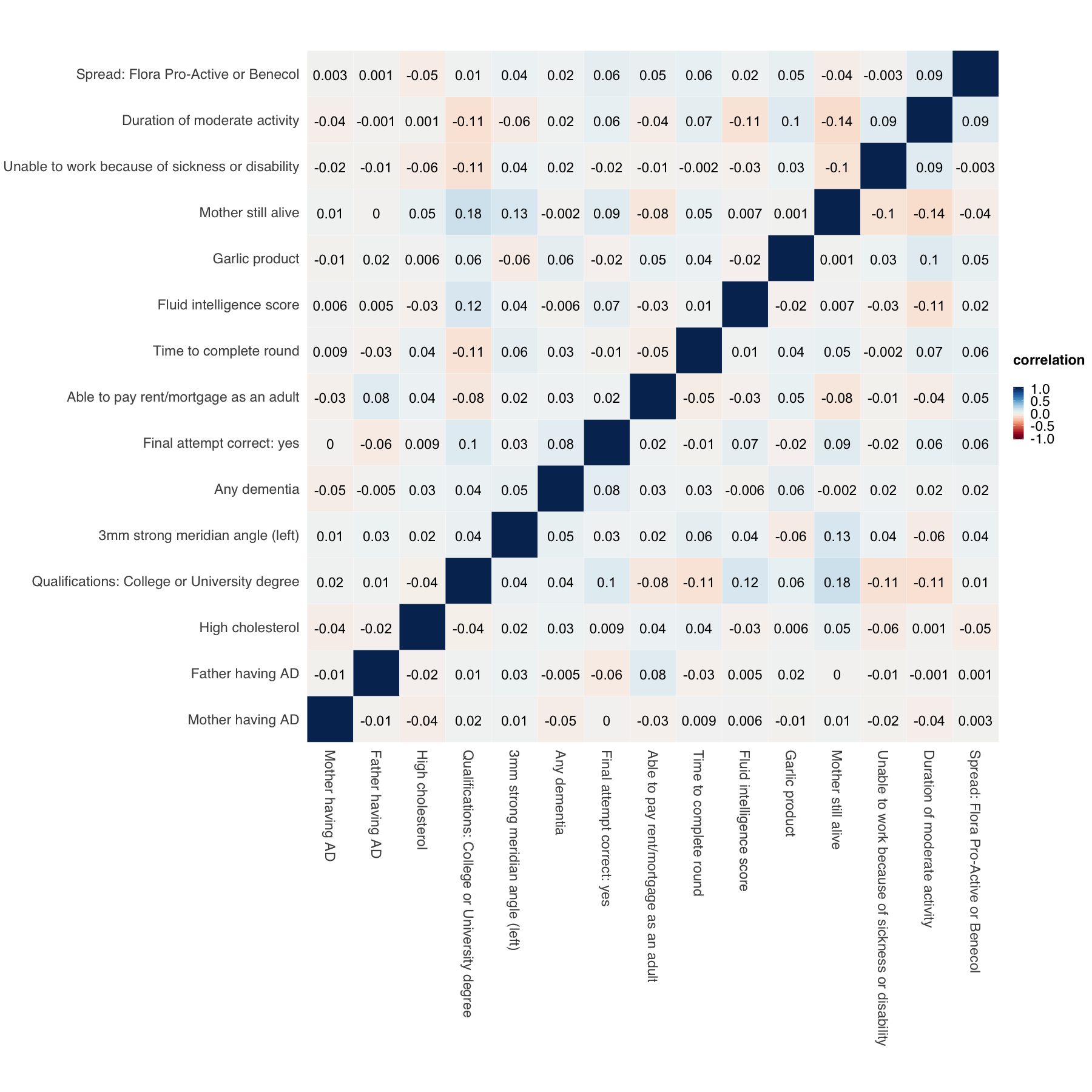

**Supplementary Figure 9. Influence of a wider *APOE* region on PRS-AD associations.** A region of more than 2Mb was removed from GWAS summary statistics for this analysis (chr19: 44,409,039-46,412,650). The horizontal and vertical axes denote association p-values before and after removal of the extended *APOE* region, respectively. Original p-values (i.e. the x-axis) were truncated at 1e-20 for visualization purposes.

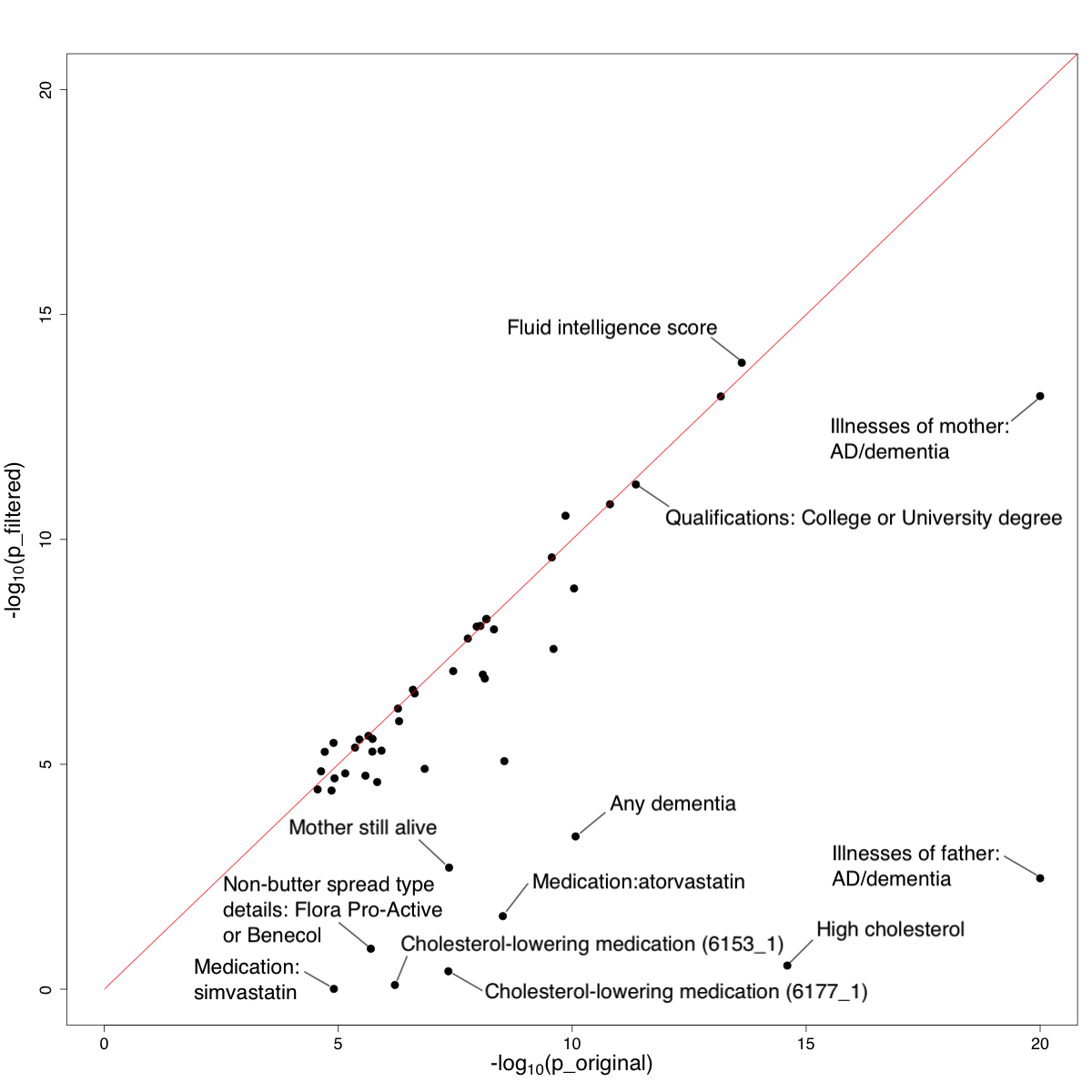

**Supplementary Figure 10. Association directions between identified AD risk factors and AD endophenotypes**. Z-scores are truncated at 5 and -5 for visualization purpose.

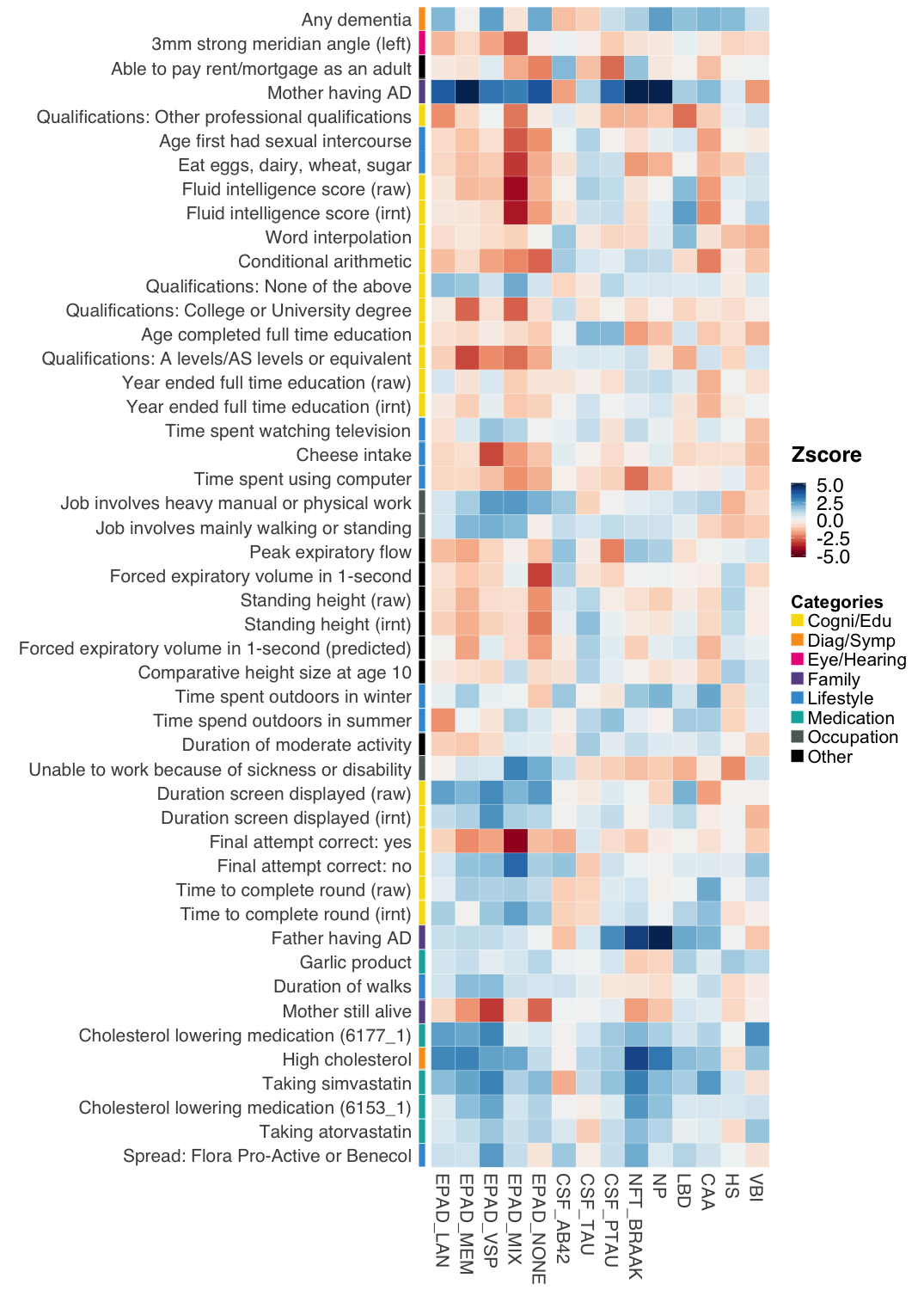

**Supplementary Figure 11. Association results for the complete set of 13 neuropathologic features for AD and other dementias. (A)** association p-values; **(B)** Association z-scores. P-values are truncated at 1e-5 and z-scores are truncated at 5 and -5 for visualization purposes.

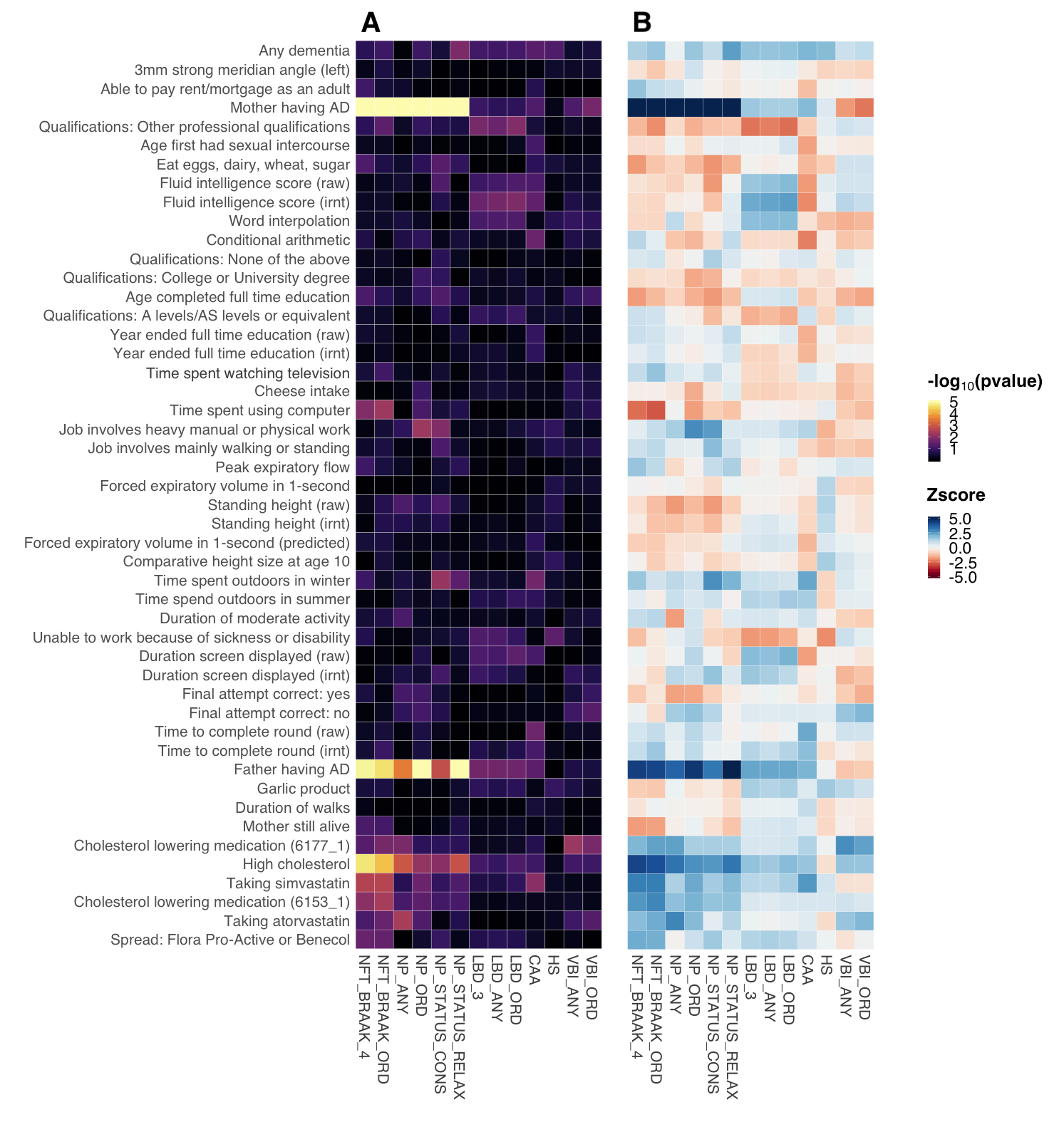

**Supplementary Tables**

**Supplementary Table 1. Type-I error rates in simulation settings 1 and 2.** Type-I error rates were calculated as the fraction of traits with p-values below the alpha value.

| **Alpha** | **Setting 1** | **Setting 2** |
| --- | --- | --- |
| 0.1 | 0.124 | 0.117 |
| 0.05 | 0.064 | 0.064 |
| 0.01 | 0.009 | 0.013 |
| 0.005 | 0.003 | 0.006 |
| 0.001 | 0.002 | 0.000 |

**Supplementary Table 2.** The list of simulation scenarios where BADGERS and regression analysis made different statistical decisions (i.e. only one method claimed statistical significance at p<0.05) in simulation setting 3.

| **Pvalue**  **cutoff** | **Trait_ID** | **reg-effect** | **reg-se** | **reg-z** | **reg-p** | **badgers-effect** | **badgers-se** | **badgers-z** | **badgers-p** |
| --- | --- | --- | --- | --- | --- | --- | --- | --- | --- |
| 0.015 | C67 | 0.614 | 0.315 | 1.947 | 0.052 | 1.535 | 1.460E-04 | 2.091 | 0.037 |
| 0.015 | 22620_1 | 0.045 | 0.024 | 1.903 | 0.057 | 0.054 | 0.029 | 2.098 | 0.036 |
| 0.015 | 20003_1140879778 | 0.393 | 0.204 | 1.929 | 0.054 | 0.398 | 4.121E-04 | 1.973 | 0.049 |
| 0.015 | 20541 | 0.060 | 0.030 | 1.985 | 0.047 | 0.057 | 0.018 | 1.942 | 0.052 |
| 0.010 | 1150_3 | 0.138 | 0.078 | 1.776 | 0.076 | 0.163 | 0.003 | 1.984 | 0.047 |
| 0.008 | 41248_1001 | 0.215 | 0.113 | 1.906 | 0.057 | 0.242 | 0.001 | 2.021 | 0.043 |
| 0.008 | 20126_3 | 0.080 | 0.043 | 1.867 | 0.062 | 0.107 | 0.008 | 2.158 | 0.031 |
| 0.008 | 3799 | 0.034 | 0.019 | 1.842 | 0.066 | 0.044 | 0.043 | 2.093 | 0.036 |
| 0.008 | 20426 | 0.029 | 0.014 | 2.098 | 0.036 | 0.023 | 0.089 | 1.784 | 0.074 |
| 0.005 | 5133_raw | 0.009 | 0.005 | 1.901 | 0.057 | 0.010 | 0.690 | 2.067 | 0.039 |
| 0.005 | 22617_2132 | 0.122 | 0.058 | 2.109 | 0.035 | 0.005 | 0.051 | 0.106 | 0.051 |
| 0.005 | 20003_1140911698 | 2.934 | 1.415 | 2.073 | 0.038 | 2.569 | 8.596E-06 | 1.921 | 0.055 |

**Supplementary Table 5. Marginal and conditional associations of 15 representative traits with AD.**

|  | **Marginal Analysis** | | | **Conditional Analysis** | | |
| --- | --- | --- | --- | --- | --- | --- |
| **Trait** | **Effect** | **SE** | **P** | **Effect** | **SE** | **P** |
| Illnesses of mother: Alzheimer's disease/dementia | 1.885 | 0.101 | 3.7e-77 | 1.808 | 0.102 | 8.1e-71 |
| Illnesses of father: Alzheimer's disease/dementia | 1.495 | 0.136 | 5.2e-28 | 1.394 | 0.138 | 3.8e-24 |
| Non-cancer illness code, self-reported: high cholesterol | 0.536 | 0.068 | 2.5e-15 | 0.535 | 0.069 | 5.3e-15 |
| Qualifications: College or University degree | -0.154 | 0.022 | 4.4e-12 | -0.137 | 0.024 | 9.9e-09 |
| 3mm strong meridian angle (left) | -0.042 | 0.010 | 2.8e-05 | -0.064 | 0.011 | 1.1e-08 |
| Any dementia | 9.084 | 1.399 | 8.5e-11 | 8.073 | 1.423 | 1.4e-08 |
| Final attempt correct: yes | -0.561 | 0.087 | 9.1e-11 | -0.494 | 0.090 | 3.8e-08 |
| Able to pay rent/mortgage as an adult | -0.086 | 0.020 | 2.3e-05 | -0.103 | 0.023 | 5.8e-06 |
| Time to complete round | 0.001 | 0.0002 | 2.8e-09 | 0.001 | 0.0002 | 7.5e-06 |
| Fluid intelligence score | -0.073 | 0.010 | 2.4e-14 | -0.043 | 0.010 | 1.3e-05 |
| Treatment/medication code: garlic product | 1.221 | 0.256 | 1.9e-06 | 1.088 | 0.253 | 1.7e-05 |
| Mother still alive | -0.408 | 0.074 | 4.3e-08 | -0.319 | 0.076 | 2.7e-05 |
| Unable to work because of sickness or disability | 0.701 | 0.124 | 1.7e-08 | 0.455 | 0.125 | 2.6e-04 |
| Duration of moderate activity | 0.146 | 0.025 | 7.4e-09 | 0.092 | 0.026 | 3.6e-04 |
| Non-butter spread type details: Flora Pro-Active or Benecol | 0.310 | 0.065 | 2.0e-06 | 0.179 | 0.073 | 0.014 |

**Supplementary Table 7. Results for Mendelian randomization based on the MR-IVW approach.** Among 1,738 heritable UK-biobank traits, 9 traits reached Bonferroni-corrected statistical significance in the meta-analysis.

| **Trait** | **ID** | **Effect** | **SE** | **P** |
| --- | --- | --- | --- | --- |
| Illnesses of mother: Alzheimer's disease/dementia | 20110_10 | 27.974 | 0.857 | 1.09E-233 |
| Illnesses of father: Alzheimer's disease/dementia | 20107_10 | 45.725 | 2.595 | 1.68E-69 |
| Any dementia | KRA_PSY_DEMENTIA | 845.495 | 161.889 | 1.76E-07 |
| Treatment/medication code: insulin product | 20003_1140883066 | -11.032 | 2.213 | 6.16E-07 |
| Non-cancer illness code, self-reported: polymyalgia rheumatica | 20002_1377 | -29.634 | 6.165 | 1.54E-06 |
| Non-cancer illness code, self-reported: high cholesterol | 20002_1473 | 10.220 | 2.220 | 4.14E-06 |
| Medication for cholesterol, blood pressure, diabetes, or take exogenous hormones: Cholesterol lowering medication | 6153_1 | 10.258 | 2.230 | 4.23E-06 |
| Medication for cholesterol, blood pressure or diabetes: Cholesterol lowering medication | 6177_1 | 6.971 | 1.646 | 2.28E-05 |
| Non-butter spread type details: Flora Pro-Active or Benecol | 2654_2 | 16.374 | 3.902 | 2.72E-05 |

**Supplementary Table 8. Results for Mendelian randomization.**

| **Trait** | **ID** | **Effect** | **SE** | **P** |
| --- | --- | --- | --- | --- |
| Illnesses of mother: Alzheimer's disease/dementia | 20110_10 | 27.974 | 0.857 | 1.09E-233 |
| Illnesses of father: Alzheimer's disease/dementia | 20107_10 | 45.725 | 2.595 | 1.68E-69 |
| Any dementia | KRA_PSY_DEMENTIA | 336.398 | 68.524 | 9.14E-07 |
| Non-cancer illness code, self-reported: high cholesterol | 20002_1473 | 10.220 | 2.220 | 4.14E-06 |
| Cholesterol lowering medication | 6153_1 | 10.258 | 2.230 | 4.23E-06 |
| Cholesterol lowering medication | 6177_1 | 6.971 | 1.646 | 2.28E-05 |
| Non-butter spread type details: Flora Pro-Active or Benecol | 2654_2 | 16.374 | 3.902 | 2.72E-05 |
| Treatment/medication code: simvastatin | 20003_1140861958 | 14.765 | 3.676 | 5.91E-05 |
| Treatment/medication code: atorvastatin | 20003_1141146234 | 28.482 | 7.293 | 9.41E-05 |
| Qualifications: A levels/AS levels or equivalent | 6138_2 | -1.473 | 0.392 | 1.71E-04 |
| Time spent watching television (TV) | 1070 | 0.828 | 0.225 | 2.38E-04 |
| Qualifications: None of the above | 6138_100 | 1.558 | 0.424 | 2.40E-04 |
| Mother still alive | 1835 | -14.999 | 4.395 | 6.44E-04 |
| Job involves mainly walking or standing | 806 | 0.609 | 0.186 | 1.07E-03 |
| Duration screen displayed | 4290_irnt | 0.438 | 0.144 | 2.35E-03 |
| FI3 : word interpolation | 4957 | -1.143 | 0.389 | 3.29E-03 |
| Qualifications: College or University degree | 6138_1 | -0.892 | 0.323 | 5.81E-03 |
| Job involves heavy manual or physical work | 816 | 0.458 | 0.178 | 0.010 |
| Forced expiratory volume in 1-second (FEV1), predicted | 20153_raw | -0.577 | 0.228 | 0.011 |
| Age completed full time education | 845 | -0.483 | 0.197 | 0.014 |
| Time spent using computer | 1080 | -0.652 | 0.267 | 0.015 |
| Time to complete round | 400_raw | 0.027 | 0.014 | 0.048 |
| Duration of walks | 874_raw | 0.006 | 0.003 | 0.050 |
| Standing height | 50_raw | -0.017 | 0.009 | 0.052 |
| Comparative height size at age 10 | 1697 | -0.220 | 0.113 | 0.052 |
| Standing height | 50_irnt | -0.148 | 0.077 | 0.055 |
| FI6 : conditional arithmetic | 4990 | -0.565 | 0.300 | 0.060 |
| Fluid intelligence score | 20016_raw | -0.129 | 0.069 | 0.062 |
| Year ended full time education | 22501_raw | -0.074 | 0.043 | 0.082 |
| Fluid intelligence score | 20016_irnt | -0.252 | 0.148 | 0.089 |
| Treatment/medication code: garlic product | 20003_1140911732 | 4.091 | 2.860 | 0.153 |
| Age first had sexual intercourse | 2139_irnt | -0.233 | 0.166 | 0.162 |
| Unable to work because of sickness or disability | 6142_4 | 1.843 | 1.361 | 0.176 |
| Time to complete round | 400_irnt | 1.896 | 1.434 | 0.186 |
| Peak expiratory flow (PEF) | 3064_raw | 0.002 | 0.001 | 0.190 |
| Time spend outdoors in summer | 1050 | 0.304 | 0.251 | 0.225 |
| Year ended full time education | 22501_irnt | -0.398 | 0.334 | 0.233 |
| Qualifications: Other professional qualifications eg: nursing, teaching | 6138_6 | -0.638 | 0.541 | 0.239 |
| Duration screen displayed | 4290_raw | 0.001 | 0.001 | 0.255 |
| Time spent outdoors in winter | 1060 | 0.467 | 0.416 | 0.262 |
| Final attempt correct: no | 4294_0 | 0.856 | 0.804 | 0.287 |
| Able to pay rent/mortgage as an adult | 20525 | -0.197 | 0.186 | 0.290 |
| 3mm strong meridian angle (left) | 5104_irnt | 0.033 | 0.121 | 0.788 |
| Never eat eggs, dairy, wheat, sugar: I eat all of the above | 6144_5 | -0.157 | 0.629 | 0.803 |
| Cheese intake | 1408 | 0.046 | 0.191 | 0.808 |
| Forced expiratory volume in 1-second (FEV1) | 3063_raw | 0.042 | 0.185 | 0.821 |
| Duration of moderate activity | 894 | 0.042 | 0.259 | 0.870 |
| Final attempt correct: yes | 4294_1 | 0.022 | 0.827 | 0.979 |

**Supplementary Table 9.** **GSMR results on the IGAP GWAS dataset.** Among 1,738 heritable UK-biobank traits, 18 traits reached Bonferroni-corrected statistical significance.

| **Trait** | **ID** | **Effect** | **SE** | **P** |
| --- | --- | --- | --- | --- |
| Immature reticulocyte fraction | 30280_irnt | 0.285 | 0.002 | 0 |
| Illnesses of mother: Alzheimer's disease/dementia | 20110_10 | 29.210 | 0.797 | 2.38E-294 |
| Heel quantitative ultrasound index (QUI), direct entry | 3147_irnt | 0.371 | 0.019 | 4.64E-82 |
| Arm fat mass (right) | 23120_raw | 1.291 | 0.103 | 2.68E-36 |
| Arm fat percentage (left) | 23123_irnt | 1.408 | 0.140 | 1.04E-23 |
| Ankle spacing width | 3143_irnt | -0.471 | 0.060 | 4.57E-15 |
| Leg fat mass (left) | 23116_irnt | -0.416 | 0.066 | 2.80E-10 |
| Standing height | 50_irnt | -0.165 | 0.029 | 1.36E-08 |
| Standing height | 50_raw | -0.017 | 0.003 | 5.71E-08 |
| Mean reticulocyte volume | 30260_irnt | -0.166 | 0.032 | 1.66E-07 |
| Leg fat percentage (right) | 23111_raw | -0.056 | 0.011 | 2.12E-07 |
| Lymphocyte percentage | 30180_raw | 0.021 | 0.004 | 1.76E-06 |
| Qualifications: A levels/AS levels or equivalent | 6138_2 | -1.515 | 0.328 | 3.77E-06 |
| Comparative height size at age 10 | 1697 | -0.229 | 0.050 | 3.77E-06 |
| Qualifications: College or University degree | 6138_1 | -0.652 | 0.143 | 4.84E-06 |
| Trunk fat-free mass | 23129_irnt | -0.231 | 0.051 | 6.20E-06 |
| Whole body fat-free mass | 23101_raw | -0.022 | 0.005 | 2.17E-05 |
| Trunk predicted mass | 23130_irnt | -0.214 | 0.051 | 2.23E-05 |

**Supplementary Table 10.** **Mendelian randomization result after removing the *APOE* region**.

| **Trait** | **ID** | **Effect** | **SE** | **P** |
| --- | --- | --- | --- | --- |
| Illnesses of mother: Alzheimer's disease/dementia | 20110_10 | 10.004 | 2.086 | 1.63E-06 |
| Qualifications: A levels/AS levels or equivalent | 6138_2 | -1.473 | 0.392 | 1.71E-04 |
| Time spent watching television (TV) | 1070 | 0.828 | 0.225 | 2.38E-04 |
| Qualifications: None of the above | 6138_100 | 1.558 | 0.424 | 2.40E-04 |
| Job involves mainly walking or standing | 806 | 0.609 | 0.186 | 0.001 |
| Duration screen displayed | 4290_irnt | 0.438 | 0.144 | 0.002 |
| FI3 : word interpolation | 4957 | -1.143 | 0.389 | 0.003 |
| Qualifications: College or University degree | 6138_1 | -0.892 | 0.323 | 0.006 |
| Job involves heavy manual or physical work | 816 | 0.458 | 0.178 | 0.010 |
| Forced expiratory volume in 1-second (FEV1), predicted | 20153_raw | -0.577 | 0.228 | 0.011 |
| Age completed full time education | 845 | -0.483 | 0.197 | 0.014 |
| Time spent using computer | 1080 | -0.652 | 0.267 | 0.015 |
| Duration of walks | 874_raw | 0.006 | 0.003 | 0.050 |
| Time to complete round | 400_raw | 0.004 | 0.002 | 0.051 |
| Standing height | 50_raw | -0.017 | 0.009 | 0.052 |
| Illnesses of father: Alzheimer's disease/dementia | 20107_10 | 5.246 | 2.701 | 0.052 |
| Comparative height size at age 10 | 1697 | -0.220 | 0.113 | 0.052 |
| Standing height | 50_irnt | -0.148 | 0.077 | 0.055 |
| FI6 : conditional arithmetic | 4990 | -0.565 | 0.300 | 0.060 |
| Fluid intelligence score | 20016_raw | -0.129 | 0.069 | 0.062 |
| Year ended full time education | 22501_raw | -0.074 | 0.043 | 0.082 |
| Fluid intelligence score | 20016_irnt | -0.252 | 0.148 | 0.089 |
| Mother still alive | 1835 | 1.223 | 0.850 | 0.150 |
| Qualifications: Other professional qualifications eg: nursing, teaching | 6138_6 | -0.768 | 0.536 | 0.151 |
| Treatment/medication code: garlic product | 20003_1140911732 | 4.091 | 2.860 | 0.153 |
| Age first had sexual intercourse | 2139_irnt | -0.233 | 0.166 | 0.162 |
| Unable to work because of sickness or disability | 6142_4 | 1.843 | 1.361 | 0.176 |
| Peak expiratory flow (PEF) | 3064_raw | 0.002 | 0.001 | 0.190 |
| Any dementia | KRA_PSY_DEMENTIA | 13.507 | 11.109 | 0.224 |
| Time spend outdoors in summer | 1050 | 0.304 | 0.251 | 0.225 |
| Year ended full time education | 22501_irnt | -0.398 | 0.334 | 0.233 |
| Duration screen displayed | 4290_raw | 0.001 | 0.001 | 0.255 |
| Time spent outdoors in winter | 1060 | 0.467 | 0.416 | 0.262 |
| Final attempt correct: no | 4294_0 | 0.856 | 0.804 | 0.287 |
| Able to pay rent/mortgage as an adult | 20525 | -0.197 | 0.186 | 0.290 |
| Cholesterol lowering medication | 6177_1 | 0.188 | 0.259 | 0.469 |
| Non-butter spread type details: Flora Pro-Active or Benecol | 2654_2 | -0.361 | 0.587 | 0.538 |
| Treatment/medication code: simvastatin | 20003_1140861958 | -0.335 | 0.600 | 0.577 |
| Time to complete round | 400_irnt | 0.098 | 0.221 | 0.659 |
| Cholesterol lowering medication | 6153_1 | -0.132 | 0.323 | 0.682 |
| Non-cancer illness code, self-reported: high cholesterol | 20002_1473 | 0.106 | 0.298 | 0.722 |
| 3mm strong meridian angle (left) | 5104_irnt | 0.033 | 0.121 | 0.788 |
| Never eat eggs, dairy, wheat, sugar: I eat all of the above | 6144_5 | -0.157 | 0.629 | 0.803 |
| Cheese intake | 1408 | 0.046 | 0.191 | 0.808 |
| Forced expiratory volume in 1-second (FEV1) | 3063_raw | 0.042 | 0.185 | 0.821 |
| Duration of moderate activity | 894 | 0.042 | 0.259 | 0.870 |
| Final attempt correct: yes | 4294_1 | 0.022 | 0.827 | 0.979 |
| Treatment/medication code: atorvastatin | 20003_1141146234 | -0.022 | 1.023 | 0.983 |

**Supplementary Table 11. Details on GWAS datasets for AD endophenotypes.**

| **Category** | **Trait** | **N** | **Ref.** |
| --- | --- | --- | --- |
| AD Subgroup | EPAD_language | 3,768 | [1, 2] |
|  | EPAD_memory | 4,097 |  |
|  | EPAD_visuospatial | 3,738 |  |
|  | EPAD_mix | 3,585 |  |
|  | EPAD_none | 4,418 |  |
| CSF Biomarkers | Ab42 | 3,115 | [3, 4] |
|  | Ab42_male | 1,515 |  |
|  | Ab42_female | 1,501 |  |
|  | Ptau | 2,833 |  |
|  | Ptau_male | 1,452 |  |
|  | Ptau_female | 1,381 |  |
|  | Tau | 3,108 |  |
|  | Tau_male | 1,508 |  |
|  | Tau_female | 1,498 |  |
| Neuropathologic Features | CAA | 2,807 | [5] |
|  | NFT_Braak_4 | 4,735 |  |
|  | NFT_Braak_ordinal | 4,707 |  |
|  | LBD_3 | 3,526 |  |
|  | LBD_any | 3,526 |  |
|  | LBD_ordinal | 3,525 |  |
|  | HS | 2,886 |  |
|  | NP_any | 4,046 |  |
|  | NP_ordinal | 4,232 |  |
|  | NP_complete | 3,702 |  |
|  | NP_primary | 4,914 |  |
|  | VBI_any | 2,764 |  |
|  | VBI_ordinal | 2,940 |  |

**Supplementary Table 19. Heritability of 48 identified traits in the marginal association analysis.** All identified traits held heritability over 0.0025

| **Trait** | **ID** | **H2** |
| --- | --- | --- |
| Standing height | 50_raw | 0.427 |
| Standing height | 50_irnt | 0.423 |
| Forced expiratory volume in 1-second (FEV1), predicted | 20153_raw | 0.422 |
| Fluid intelligence score | 20016_raw | 0.228 |
| Comparative height size at age 10 | 1697 | 0.222 |
| Fluid intelligence score | 20016_irnt | 0.221 |
| Forced expiratory volume in 1-second (FEV1) | 3063_raw | 0.178 |
| Qualifications: College or University degree | 6138_1 | 0.162 |
| Age first had sexual intercourse | 2139_irnt | 0.154 |
| Year ended full time education | 22501_raw | 0.148 |
| Year ended full time education | 22501_irnt | 0.136 |
| Duration screen displayed | 4290_irnt | 0.113 |
| Age completed full time education | 845 | 0.104 |
| Time spent watching television (TV) | 1070 | 0.104 |
| Peak expiratory flow (PEF) | 3064_raw | 0.103 |
| Qualifications: None of the above | 6138_100 | 0.103 |
| Time spent using computer | 1080 | 0.096 |
| Qualifications: A levels/AS levels or equivalent | 6138_2 | 0.091 |
| FI6: conditional arithmetic | 4990 | 0.091 |
| Time to complete round | 400_irnt | 0.087 |
| FI3: word interpolation | 4957 | 0.084 |
| Job involves heavy manual or physical work | 816 | 0.081 |
| Medication for cholesterol, blood pressure or diabetes: Cholesterol lowering medication | 6177_1 | 0.080 |
| 3mm strong meridian angle (left) | 5104_irnt | 0.077 |
| Job involves mainly walking or standing | 806 | 0.076 |
| Time spend outdoors in summer | 1050 | 0.072 |
| Cheese intake | 1408 | 0.068 |
| Time to complete round | 400_raw | 0.062 |
| Duration screen displayed | 4290_raw | 0.060 |
| Medication for cholesterol, blood pressure, diabetes, or take exogenous hormones: Cholesterol lowering medication | 6153_1 | 0.057 |
| Non-cancer illness code, self-reported: high cholesterol | 20002_1473 | 0.047 |
| Never eat eggs, dairy, wheat, sugar: I eat all of the above | 6144_5 | 0.047 |
| Time spent outdoors in winter | 1060 | 0.044 |
| Qualifications: Other professional qualifications eg: nursing, teaching | 6138_6 | 0.044 |
| Duration of walks | 874_raw | 0.040 |
| Duration of moderate activity | 894 | 0.032 |
| Treatment/medication code: simvastatin | 20003_1140861958 | 0.031 |
| Current employment status: Unable to work because of sickness or disability | 6142_4 | 0.024 |
| Able to pay rent/mortgage as an adult | 20525 | 0.023 |
| Final attempt correct: yes | 4294_1 | 0.021 |
| Final attempt correct: no | 4294_0 | 0.021 |
| Illnesses of mother: Alzheimer's disease/dementia | 20110_10 | 0.018 |
| Treatment/medication code: atorvastatin | 20003_1141146234 | 0.014 |
| Illnesses of father: Alzheimer's disease/dementia | 20107_10 | 0.014 |
| Non-butter spread type details: Flora Pro-Active or Benecol | 2654_2 | 0.014 |
| Mother still alive | 1835 | 0.005 |
| Any dementia | KRA_PSY_DEMENTIA | 0.003 |
| Treatment/medication code: garlic product | 20003_1140911732 | 0.003 |

**Acknowledgements to ADGC**

The National Institutes of Health, National Institute on Aging (NIH-NIA) supported this work through the following grants: ADGC, U01 AG032984, RC2 AG036528; Samples from the National Cell Repository for Alzheimer’s Disease (NCRAD), which receives government support under a cooperative agreement grant (U24 AG21886) awarded by the National Institute on Aging (NIA), were used in this study. We thank contributors who collected samples used in this study, as well as patients and their families, whose help and participation made this work possible; Data for this study were prepared, archived, and distributed by the National Institute on Aging Alzheimer’s Disease Data Storage Site (NIAGADS) at the University of Pennsylvania (U24-AG041689-01); NACC, U01 AG016976; NIA LOAD (Columbia University), U24 AG026395, U24 AG026390, R01AG041797; Banner Sun Health Research Institute P30 AG019610; Boston University, P30 AG013846, U01 AG10483, R01 CA129769, R01 MH080295, R01 AG017173, R01 AG025259, R01 AG048927, R01AG33193, R01 AG009029; Columbia University, P50 AG008702, R37 AG015473, R01 AG037212, R01 AG028786; Duke University, P30 AG028377, AG05128; Emory University, AG025688; Group Health Research Institute, UO1 AG006781, UO1 HG004610, UO1 HG006375, U01 HG008657; Indiana University, P30 AG10133, R01 AG009956, RC2 AG036650; Johns Hopkins University, P50 AG005146, R01 AG020688; Massachusetts General Hospital, P50 AG005134; Mayo Clinic, P50 AG016574, R01 AG032990, KL2 RR024151; Mount Sinai School of Medicine, P50 AG005138, P01 AG002219; New York University, P30 AG08051, UL1 RR029893, 5R01AG012101, 5R01AG022374, 5R01AG013616, 1RC2AG036502, 1R01AG035137; North Carolina A&T University, P20 MD000546, R01 AG28786-01A1; Northwestern University, P30 AG013854; Oregon Health & Science University, P30 AG008017, R01 AG026916; Rush University, P30 AG010161, R01 AG019085, R01 AG15819, R01 AG17917, R01 AG030146, R01 AG01101, RC2 AG036650, R01 AG22018; TGen, R01 NS059873; University of Alabama at Birmingham, P50 AG016582; University of Arizona, R01 AG031581; University of California, Davis, P30 AG010129; University of California, Irvine, P50 AG016573; University of California, Los Angeles, P50 AG016570; University of California, San Diego, P50 AG005131; University of California, San Francisco, P50 AG023501, P01 AG019724; University of Kentucky, P30 AG028383, AG05144; University of Michigan, P50 AG008671; University of Pennsylvania, P30 AG010124; University of Pittsburgh, P50 AG005133, AG030653, AG041718, AG07562, AG02365; University of Southern California, P50 AG005142; University of Texas Southwestern, P30 AG012300; University of Miami, R01 AG027944, AG010491, AG027944, AG021547, AG019757; University of Washington, P50 AG005136, R01 AG042437; University of Wisconsin, P50 AG033514; Vanderbilt University, R01 AG019085; and Washington University, P50 AG005681, P01 AG03991, P01 AG026276. The Kathleen Price Bryan Brain Bank at Duke University Medical Center is funded by NINDS grant # NS39764, NIMH MH60451 and by Glaxo Smith Kline. Support was also from the Alzheimer’s Association (LAF, IIRG-08-89720; MP-V, IIRG-05-14147), the US Department of Veterans Affairs Administration, Office of Research and Development, Biomedical Laboratory Research Program, and BrightFocus Foundation (MP-V, A2111048). P.S.G.-H. is supported by Wellcome Trust, Howard Hughes Medical Institute, and the Canadian Institute of Health Research. Genotyping of the TGEN2 cohort was supported by Kronos Science. The TGen series was also funded by NIA grant AG041232 to AJM and MJH, The Banner Alzheimer’s Foundation, The Johnnie B. Byrd Sr. Alzheimer’s Institute, the Medical Research Council, and the state of Arizona and also includes samples from the following sites: Newcastle Brain Tissue Resource (funding via the Medical Research Council, local NHS trusts and Newcastle University), MRC London Brain Bank for Neurodegenerative Diseases (funding via the Medical Research Council),South West Dementia Brain Bank (funding via numerous sources including the Higher Education Funding Council for England (HEFCE), Alzheimer’s Research Trust (ART), BRACE as well as North Bristol NHS Trust Research and Innovation Department and DeNDRoN), The Netherlands Brain Bank (funding via numerous sources including Stichting MS Research, Brain Net Europe, Hersenstichting Nederland Breinbrekend Werk, International Parkinson Fonds, Internationale Stiching Alzheimer Onderzoek), Institut de Neuropatologia, Servei Anatomia Patologica, Universitat de Barcelona. ADNI data collection and sharing was funded by the National Institutes of Health Grant U01 AG024904 and Department of Defense award number W81XWH-12-2-0012. ADNI is funded by the National Institute on Aging, the National Institute of Biomedical Imaging and Bioengineering, and through generous contributions from the following: AbbVie, Alzheimer’s Association; Alzheimer’s Drug Discovery Foundation; Araclon Biotech; BioClinica, Inc.; Biogen; Bristol-Myers Squibb Company; CereSpir, Inc.; Eisai Inc.; Elan Pharmaceuticals, Inc.; Eli Lilly and Company; EuroImmun; F. Hoffmann-La Roche Ltd and its affiliated company Genentech, Inc.; Fujirebio; GE Healthcare; IXICO Ltd.; Janssen Alzheimer Immunotherapy Research & Development, LLC.; Johnson & Johnson Pharmaceutical Research & Development LLC.; Lumosity; Lundbeck; Merck & Co., Inc.; Meso Scale Diagnostics, LLC.; NeuroRx Research; Neurotrack Technologies; Novartis Pharmaceuticals Corporation; Pfizer Inc.; Piramal Imaging; Servier; Takeda Pharmaceutical Company; and Transition Therapeutics. The Canadian Institutes of Health Research is providing funds to support ADNI clinical sites in Canada. Private sector contributions are facilitated by the Foundation for the National Institutes of Health (www.fnih.org). The grantee organization is the Northern California Institute for Research and Education, and the study is coordinated by the Alzheimer's Disease Cooperative Study at the University of California, San Diego. ADNI data are disseminated by the Laboratory for Neuro Imaging at the University of Southern California. We thank Drs. D. Stephen Snyder and Marilyn Miller from NIA who are *ex-officio* ADGC members.

We would also like to thank all the members of ADGC: Erin Abner ^1^, Perrie M. Adams ^2^, Marilyn S. Albert ^3^, Roger L. Albin ^4-,6^, Liana G. Apostolova ^7-10^, Steven E. Arnold ^11^, Sanjay Asthana ^12-14^, Craig S. Atwood ^12-14^, Clinton T. Baldwin ^15^, Robert C. Barber ^16^, Lisa L. Barnes ^17-19^, Sandra Barral ^20-22^, Thomas G. Beach ^23^, James T. Becker ^24^, Gary W. Beecham ^25,26^, Duane Beekly ^27^, David A. Bennett ^17,19^, Eileen H. Bigio ^28,29^, Thomas D. Bird ^30,31^, Deborah Blacker ^32,33^, Bradley F. Boeve ^34^, James D. Bowen ^35^, Adam Boxer ^36^, James R. Burke ^37^, Jeffrey M. Burns ^38^, Joseph D. Buxbaum ^39-41^, Nigel J. Cairns ^42^, Laura B. Cantwell ^43^, Chuanhai Cao ^44^, Chris S. Carlson ^45^, Cynthia M. Carlsson ^12-14^, Regina M. Carney ^46^, Minerva M. Carrasquillo ^47^, Helena C. Chui ^48^, Paul K. Crane ^49^, David H. Cribbs ^50^, Elizabeth A. Crocco ^46^, Carlos Cruchaga ^51^, Philip L. De Jager ^52,53^, Charles DeCarli ^54^, Malcolm Dick ^55^, Dennis W. Dickson ^47^, Rachelle S. Doody ^56^, Ranjan Duara ^57^, Nilufer Ertekin-Taner ^47,58^, Denis A. Evans ^59^, Kelley M. Faber ^8^, Thomas J. Fairchild ^60^, Kenneth B. Fallon ^61^, David W. Fardo ^62^, Martin R. Farlow ^63^, Lindsay A. Farrer ^64-68^, Steven Ferris ^69^, Tatiana M. Foroud ^8^, Matthew P. Frosch ^70^, Douglas R. Galasko ^71^, Marla Gearing ^72,73^, Daniel H. Geschwind ^74^, Bernardino Ghetti ^75^, John R. Gilbert ^25,26^, Alison M. Goate ^39^, Neill R. Graff-Radford ^47,58^, Robert C. Green ^76^, John H. Growdon ^77^, Jonathan L. Haines ^78^, Hakon Hakonarson ^79^, Ronald L. Hamilton ^80^, Kara L. Hamilton-Nelson ^25^, John Hardy ^81^, Lindy E. Harrell ^82^, Lawrence S. Honig ^20^, Ryan M. Huebinger ^83^, Matthew J. Huentelman ^84^, Christine M. Hulette ^85^, Bradley T. Hyman ^77^, Gail P. Jarvik ^86,87^, Lee-Way Jin ^88^, Gyungah Jun ^15,64,68^, M. Ilyas Kamboh ^89,90^, Anna Karydas ^36^, Mindy J. Katz ^91^, John S.K. Kauwe ^92^, Jeffrey A. Kaye ^93,94^, C. Dirk Keene ^95^, Ronald Kim ^96^, Neil W. Kowall^67,97^, Joel H. Kramer ^98^, Walter A. Kukull ^99^, Brian W. Kunkle ^25^, Amanda P. Kuzma ^43^, Frank M. LaFerla ^100^, James J. Lah ^101^, Eric B. Larson ^49,102^, James B. Leverenz ^103^, Allan I. Levey ^101^, Ge Li ^31,104^, Andrew P. Lieberman ^105^, Richard B. Lipton ^91^, Oscar L. Lopez ^90^, Kathryn L. Lunetta ^64^, Constantine G. Lyketsos ^106^, John Malamon ^43^, Daniel C. Marson ^82^, Eden R. Martin ^25,26^, Frank Martiniuk ^107^, Deborah C. Mash ^108^, Eliezer Masliah ^71,109^, Richard Mayeux ^20,21^, Wayne C. McCormick ^49^, Susan M. McCurry ^110^, Andrew N. McDavid ^45^, Stefan McDonough ^111^, Ann C. McKee ^67,97^, Marsel Mesulam ^29,112^, Bruce L. Miller^36^, Carol A. Miller ^113^, Joshua W. Miller ^88^, Thomas J. Montine ^95^, John C. Morris ^42,114^, Shubhabrata Mukherjee ^49^, Amanda J. Myers ^46^, Adam C. Naj ^43^, Sid O’Bryant ^115^, John M. Olichney ^54^, Joseph E. Parisi^116^, Henry L. Paulson ^117^, Margaret A. Pericak-Vance ^25,26^, Elaine Peskind ^104^, Ronald C. Petersen ^34^, Aimee Pierce ^50^, Wayne W. Poon ^55^, Huntington Potter ^118^, Liming Qu ^43^, Joseph F. Quinn ^93,94^, Ashok Raj^44^, Murray Raskind ^104^, Eric M. Reiman ^84,119-121^, Barry Reisberg ^69,122^, Joan S. Reisch ^123^, Christiane Reitz^20-22,124^, John M. Ringman ^48^, Erik D. Roberson ^82^, Ekaterina Rogaeva ^125^, Howard J. Rosen ^36^, Roger N. Rosenberg ^127^, Donald R. Royall ^128^, Mark A. Sager ^13^, Mary Sano ^40^, Andrew J. Saykin ^7,8^, Gerard D. Schellenberg ^43^, Julie A. Schneider ^17,19,129^, Lon S. Schneider ^48,130^, William W. Seeley ^36^, Amanda G. Smith^44^, Joshua A. Sonnen ^95^, Salvatore Spina ^75^, Peter St George-Hyslop ^131,132^, Robert A. Stern ^67^, Russell H. Swerdlow ^38^, Rudolph E. Tanzi ^77^, John Q. Trojanowski ^133^, Juan C. Troncoso ^134^, Debby W. Tsuang ^31,104^, Otto Valladares ^43^, Vivianna M. Van Deerlin ^133^, Linda J. Van Eldik ^135^, Badri N. Vardarajan ^20-22^, Harry V. Vinters ^136,137^, Jean Paul Vonsattel ^138^, Li-San Wang ^43^, Sandra Weintraub ^28,29^, Kathleen A. Welsh-Bohmer^37,139^, Kirk C. Wilhelmsen ^140^, Jennifer Williamson ^20^, Thomas S. Wingo ^101^, Randall L. Woltjer ^141^, Clinton B. Wright ^142^, Chuang-Kuo Wu ^143^, Steven G. Younkin ^47^, Chang-En Yu ^49^, Lei Yu ^17,19^, Yi Zhao ^43^

^1^Sanders-Brown Center on Aging, College of Public Health, Department of Epidemiology, University of Kentucky, Lexington, Kentucky, ^2^Department of Psychiatry, University of Texas Southwestern Medical Center, Dallas, Texas, ^3^Department of Neurology, Johns Hopkins University, Baltimore, Maryland, ^4^Department of Neurology, University of Michigan, Ann Arbor, Michigan, ^5^Geriatric Research, Education and Clinical Center (GRECC), VA Ann Arbor Healthcare System (VAAAHS), Ann Arbor, Michigan, ^6^Michigan Alzheimer Disease Center, Ann Arbor, Michigan, ^7^Department of Radiology, Indiana University, Indianapolis, Indiana, ^8^Department of Medical and Molecular Genetics, Indiana University, Indianapolis, Indiana, ^9^Indian Alzheimer's Disease Center, Indiana University, Indianapolis, Indiana, ^10^Department of Neurology, Indiana University, Indianapolis, Indiana, ^11^Department of Psychiatry, University of Pennsylvania Perelman School of Medicine, Philadelphia, Pennsylvania, ^12^Geriatric Research, Education and Clinical Center (GRECC), University of Wisconsin, Madison, Wisconsin, ^13^Department of Medicine, University of Wisconsin, Madison, Wisconsin, ^14^Wisconsin Alzheimer's Disease Research Center, Madison, Wisconsin, ^1^ Department of Medicine (Genetics Program), Boston University, Boston, Massachusetts, ^16^Department of Pharmacology and Neuroscience, University of North Texas Health Science Center, Fort Worth, Texas, ^17^Department of Neurological Sciences, Rush University Medical Center, Chicago, Illinois, ^18^Department of Behavioral Sciences, Rush University Medical Center, Chicago, Illinois, ^19^Rush Alzheimer's Disease Center, Rush University Medical Center, Chicago, Illinois, ^20^Taub Institute on Alzheimer's Disease and the Aging Brain, Department of Neurology, Columbia University, New York, New York, ^21^Gertrude H. Sergievsky Center, Columbia University, New York, New York, ^22^Department of Neurology, Columbia University, New York, New York, ^23^Civin Laboratory for Neuropathology, Banner Sun Health Research Institute, Phoenix, Arizona, ^24^Departments of Psychiatry, Neurology, and Psychology, University of Pittsburgh School of Medicine, Pittsburgh, Pennsylvania, ^25^The John P. Hussman Institute for Human Genomics, University of Miami, Miami, Florida, ^26^Dr. John T. Macdonald Foundation Department of Human Genetics, University of Miami, Miami, Florida, ^27^National Alzheimer's Coordinating Center, University of Washington, Seattle, Washington, ^28^Department of Pathology, Northwestern University Feinberg School of Medicine, Chicago, Illinois, ^29^Cognitive Neurology and Alzheimer's Disease Center, Northwestern University Feinberg School of Medicine, Chicago, Illinois, ^30^Department of Neurology, University of Washington, Seattle, Washington, ^31^VA Puget Sound Health Care System/GRECC, Seattle, Washington, ^32^Department of Epidemiology, Harvard School of Public Health, Boston, Massachusetts, ^33^Department of Psychiatry, Massachusetts General Hospital/Harvard Medical School, Boston, Massachusetts, ^34^Department of Neurology, Mayo Clinic, Rochester, Minnesota, ^35^Swedish Medical Center, Seattle, Washington, ^36^Department of Neurology, University of California San Francisco, San Francisco, California, ^37^Department of Medicine, Duke University, Durham, North Carolina, ^38^University of Kansas Alzheimer’s Disease Center, University of Kansas Medical Center, Kansas City, Kansas, ^39^Department of Neuroscience, Mount Sinai School of Medicine, New York, New York, ^40^Department of Psychiatry, Mount Sinai School of Medicine, New York, New York, ^41^Departments of Genetics and Genomic Sciences, Mount Sinai School of Medicine, New York, New York, ^42^Department of Pathology and Immunology, Washington University, St. Louis, Missouri, ^43^Penn Neurodegeneration Genomics Center, Department of Pathology and Laboratory Medicine, University of Pennsylvania Perelman School of Medicine, Philadelphia, Pennsylvania, ^44^USF Health Byrd Alzheimer's Institute, University of South Florida, Tampa, Florida, ^45^Fred Hutchinson Cancer Research Center, Seattle, Washington, ^46^Department of Psychiatry and Behavioral Sciences, Miller School of Medicine, University of Miami, Miami, Florida, ^47^Department of Neuroscience, Mayo Clinic, Jacksonville, Florida, ^48^Department of Neurology, University of Southern California, Los Angeles, California, ^49^Department of Medicine, University of Washington, Seattle, Washington, ^50^Department of Neurology, University of California Irvine, Irvine, California, ^51^Department of Psychiatry and Hope Center Program on Protein Aggregation and Neurodegeneration, Washington University School of Medicine, St. Louis, Missouri, ^52^Program in Translational NeuroPsychiatric Genomics, Institute for the Neurosciences, Department of Neurology & Psychiatry, Brigham and Women's Hospital and Harvard Medical School, Boston, Massachusetts, ^53^Program in Medical and Population Genetics, Broad Institute, Cambridge, Massachusetts, ^54^Department of Neurology, University of California Davis, Sacramento, California, ^55^Institute for Memory Impairments and Neurological Disorders, University of California Irvine, Irvine, California, ^56^Alzheimer's Disease and Memory Disorders Center, Baylor College of Medicine, Houston, Texas, ^57^Wien Center for Alzheimer's Disease and Memory Disorders, Mount Sinai Medical Center, Miami Beach, Florida, ^58^Department of Neurology, Mayo Clinic, Jacksonville, Florida, ^59^Rush Institute for Healthy Aging, Department of Internal Medicine, Rush University Medical Center, Chicago, Illinois, ^60^Office of Strategy and Measurement, University of North Texas Health Science Center, Fort Worth, Texas, ^61^Department of Pathology, University of Alabama at Birmingham, Birmingham, Alabama, ^62^Sanders-Brown Center on Aging, Department of Biostatistics, University of Kentucky, Lexington, Kentucky, ^63^Department of Neurology, Indiana University, Indianapolis, Indiana, ^64^Department of Biostatistics, Boston University, Boston, Massachusetts, ^65^Department of Epidemiology, Boston University, Boston, Massachusetts, ^66^Department of Medicine (Biomedical Genetics), Boston University, Boston, Massachusetts, ^67^Department of Neurology, Boston University, Boston, Massachusetts, ^68^Department of Ophthalmology, Boston University, Boston, Massachusetts, ^69^Department of Psychiatry, New York University, New York, New York, ^70^C.S. Kubik Laboratory for Neuropathology, Massachusetts General Hospital, Charlestown, Massachusetts, ^71^Department of Neurosciences, University of California San Diego, La Jolla, California, ^72^Department of Pathology and Laboratory Medicine, Emory University, Atlanta, Georgia, ^73^Emory Alzheimer's Disease Center, Emory University, Atlanta, Georgia, ^74^Neurogenetics Program, University of California Los Angeles, Los Angeles, California, ^75^Department of Pathology and Laboratory Medicine, Indiana University, Indianapolis, Indiana, ^76^Division of Genetics, Department of Medicine and Partners Center for Personalized Genetic Medicine, Brigham and Women's Hospital and Harvard Medical School, Boston, Massachusetts, ^77^Department of Neurology, Massachusetts General Hospital/Harvard Medical School, Boston, Massachusetts, ^78^Department of Epidemiology and Biostatistics, Case Western Reserve University, Cleveland, Ohio, ^79^Center for Applied Genomics, Children's Hospital of Philadelphia, Philadelphia, Pennsylvania, ^80^Department of Pathology (Neuropathology), University of Pittsburgh, Pittsburgh, Pennsylvania, ^81^Institute of Neurology, University College London, Queen Square, London, United Kingdom, ^82^Department of Neurology, University of Alabama at Birmingham, Birmingham, Alabama, ^83^Department of Surgery, University of Texas Southwestern Medical Center, Dallas, Texas, ^84^Neurogenomics Division, Translational Genomics Research Institute, Phoenix, Arizona, ^85^Department of Pathology, Duke University, Durham, North Carolina, ^86^Department of Genome Sciences, University of Washington, Seattle, Washington, ^87^Department of Medicine (Medical Genetics), University of Washington, Seattle, Washington, ^88^Department of Pathology and Laboratory Medicine, University of California Davis, Sacramento, California, ^89^Department of Human Genetics, University of Pittsburgh, Pittsburgh, Pennsylvania, ^90^University of Pittsburgh Alzheimer's Disease Research Center, Pittsburgh, Pennsylvania, ^91^Department of Neurology, Albert Einstein College of Medicine, New York, New York, ^92^Department of Biology, Brigham Young University, Provo, Utah, ^93^Department of Neurology, Oregon Health & Science University, Portland, Oregon, ^94^Department of Neurology, Portland Veterans Affairs Medical Center, Portland, Oregon, ^95^Department of Pathology, University of Washington, Seattle, Washington, ^96^Department of Pathology and Laboratory Medicine, University of California Irvine, Irvine, California, ^97^Department of Pathology, Boston University, Boston, Massachusetts, ^98^Department of Neuropsychology, University of California San Francisco, San Francisco, California, ^99^Department of Epidemiology, University of Washington, Seattle, Washington, ^100^Department of Neurobiology and Behavior, University of California Irvine, Irvine, California, ^101^Department of Neurology, Emory University, Atlanta, Georgia, ^102^Group Health Research Institute, Group Health, Seattle, Washington, ^103^Cleveland Clinic Lou Ruvo Center for Brain Health, Cleveland Clinic, Cleveland, Ohio, ^104^Department of Psychiatry and Behavioral Sciences, University of Washington School of Medicine, Seattle, Washington, ^105^Department of Pathology, University of Michigan, Ann Arbor, Michigan, ^106^Department of Psychiatry, Johns Hopkins University, Baltimore, Maryland, ^107^Department of Medicine - Pulmonary, New York University, New York, New York, ^108^Department of Neurology, University of Miami, Miami, Florida, ^109^Department of Pathology, University of California San Diego, La Jolla, California, ^110^School of Nursing Northwest Research Group on Aging, University of Washington, Seattle, Washington, ^111^PharmaTherapeutics Clinical Research, Pfizer Worldwide Research and Development, Cambridge, Massachusetts, ^112^Department of Neurology, Northwestern University Feinberg School of Medicine, Chicago, Illinois, ^113^Department of Pathology, University of Southern California, Los Angeles, California, ^114^Department of Neurology, Washington University, St. Louis, Missouri, ^115^Internal Medicine, Division of Geriatrics, University of North Texas Health Science Center, Fort Worth, Texas, ^116^Department of Laboratory Medicine and Pathology, Mayo Clinic, Rochester, Minnesota, ^117^Michigan Alzheimer's Disease Center, Department of Neurology, University of Michigan, Ann Arbor, Michigan, ^118^Department of Neurology, University of Colorado School of Medicine, Aurora, Colorado, ^119^Arizona Alzheimer’s Consortium, Phoenix, Arizona, ^120^Banner Alzheimer's Institute, Phoenix, Arizona, ^121^Department of Psychiatry, University of Arizona, Phoenix, Arizona, ^122^Alzheimer's Disease Center, New York University, New York, New York, ^123^Department of Clinical Sciences, University of Texas Southwestern Medical Center, Dallas, Texas, ^124^Department of Epidemiology, Columbia University, New York, New York, ^125^ Tanz Centre for Research in Neurodegenerative Disease, University of Toronto, Toronto, Ontario, ^127^Department of Neurology, University of Texas Southwestern, Dallas, Texas, ^128^Departments of Psychiatry, Medicine, Family & Community Medicine, South Texas Veterans Health Administration Geriatric Research Education & Clinical Center (GRECC), UT Health Science Center at San Antonio, San Antonio, Texas, ^129^Department of Pathology (Neuropathology), Rush University Medical Center, Chicago, Illinois, ^130^Department of Psychiatry, University of Southern California, Los Angeles, California, ^131^Tanz Centre for Research in Neurodegenerative Disease, University of Toronto, Toronto, Ontario, ^132^Cambridge Institute for Medical Research and Department of Clinical Neurosciences, University of Cambridge, Cambridge, United Kingdom, ^133^Department of Pathology and Laboratory Medicine, University of Pennsylvania Perelman School of Medicine, Philadelphia, Pennsylvania, ^134^Department of Pathology, Johns Hopkins University, Baltimore, Maryland, ^135^Sanders-Brown Center on Aging, Department of Anatomy and Neurobiology, University of Kentucky, Lexington, Kentucky, ^136^Department of Neurology, University of California Los Angeles, Los Angeles, California, ^137^Department of Pathology & Laboratory Medicine, University of California Los Angeles, Los Angeles, California, ^138^Taub Institute on Alzheimer's Disease and the Aging Brain, Department of Pathology, Columbia University, New York, New York, ^139^Department of Psychiatry & Behavioral Sciences, Duke University, Durham, North Carolina, ^140^Department of Genetics, University of North Carolina Chapel Hill, Chapel Hill, North Carolina, ^141^Department of Pathology, Oregon Health & Science University, Portland, Oregon, ^142^Evelyn F. McKnight Brain Institute, Department of Neurology, Miller School of Medicine, University of Miami, Miami, Florida, ^143^Departments of Neurology, Pharmacology & Neuroscience, Texas Tech University Health Science Center, Lubbock, Texas.
